# Impact of transposable elements on methylation and gene expression across natural accessions of *Brachypodium distachyon*

**DOI:** 10.1101/2020.06.16.154047

**Authors:** Michele Wyler, Christoph Stritt, Jean-Claude Walser, Célia Baroux, Anne C. Roulin

## Abstract

Transposable elements (TEs) constitute a large fraction of plant genomes and are mostly present in a transcriptionally silent state through repressive epigenetic modifications such as DNA methylation. TE silencing is believed to influence the regulation of adjacent genes, possibly as DNA methylation spreads away from the TE. Whether this is a general principle or a context-dependent phenomenon is still under debate, pressing for studying the relationship between TEs, DNA methylation and nearby gene expression in additional plant species. Here we used the grass *Brachypodium distachyon* as a model and produced DNA methylation and transcriptome profiles for eleven natural accessions. In contrast to what is observed in *Arabidopsis thaliana*, we found that TEs have a limited impact on methylation spreading and that only few TE families are associated to a low expression of their adjacent genes. Interestingly, we found that a subset of TE insertion polymorphisms is associated with differential gene expression across accessions. Thus, although not having a global impact on gene expression, distinct TE insertions may contribute to specific gene expression patterns in *B. distachyon*.

**Significance statement:** Transposable elements (TEs) are a major component of plant genomes and a source of genetic and epigenetic innovations underlying adaptation to changing environmental conditions. Yet molecular evidence linking TE silencing and nearby gene expression are lacking for many plant species. We show that in the model grass Brachypodium DNA methylation spreads over very short distances around TEs, with an influence on gene expression for a small subset of TE families.

## Introduction

Since their discovery by Barbara McClintock in the late 1940’s (McClintock 1947), transposable elements (TEs) have been shown to populate a large fraction of eukaryotic genomes and to be an important source of genetic and phenotypic diversity. This is explained by the observation that TEs have the ability to modulate nearby gene expression and can remodel genomes when activated by stress (Grandbastien 1998; Grandbastien 2005; Thieme et al. 2017), thereby providing raw diversity necessary for adaptation to changing environments (Bonchev and Parisod 2013; Rey et al. 2016). Remarkable examples of single-trait modulation by a TE insertion providing a selective advantage have been reported in both plants and animals (Guio et al. 2014; Van’t Hof et al. 2016; Horváth et al. 2017).

While TEs can provide selective advantages, their mutagenic properties may at the same time represent a threat for the integrity of the host genome (McClintock 1950; Shang et al. 2017; Thind et al. 2018). In response, genomes have evolved several layers of defence to contain TE activity. In plants, transcriptional repression of TEs is mediated by the RNA-directed DNA methylation (RdDM) pathway (for review Sigman and Slotkin 2016; Zhang et al. 2018). Initiated by the plant-specific RNA polymerase IV (*Pol IV*), TE transcription leads to the formation of 24 nt siRNA targeting DNA methylation at the TE source locus. Acting as a double-edged sword, DNA (and histone) methylation may spread to neighbouring regions and possibly contribute the alteration of adjacent gene expression (Hollister et al. 2010; Wang et al. 2013).

Recent studies reported on the impact of TEs on DNA methylation and expression of nearby genes in different plant species (Eichten et al. 2012; Makarevitch et al. 2015; Eichten et al. 2016; Choi and Purugganan 2017; Wang et al. 2018). Yet, most of our knowledge on the regulatory mechanisms largely rely on the eudicot model *Arabidopsis thaliana* (for review Sigman and Slotkin 2016; Zhang et al. 2018). In *A. thaliana*, the presence of TEs tends to be associated with a reduced expression of adjacent genes (Wang et al. 2013), but whether this also occurs in other species remains to be demonstrated (Choi and Purugganan 2017). In this context, we contributed an analysis of DNA methylation and gene expression in the model grass *Brachypodium distachyon*.

Established as an important model for functional genomics, the small genome of *B. distachyon* (272 Mb; 30% of TEs) is fully sequenced and assembled into five chromosomes (The International Brachypodium Consortium 2010). Comparative genomics and cytogenetic analyses (Hasterok et al. 2019) showed that it underwent multiple and independent nested chromosome fusions which resulted in a higher gene density at the former distal regions of the inserted chromosomes. Furthermore, TEs are much more pervasive than in *A. thaliana*, and although their dynamics is well characterized in *B. distachyon* (Stritt et al. 2018; Stritt et al. 2019), little is know about their functional impact. We therefore asked to what extent TEs influence DNA methylation and gene expression in *B. distachyon*, and how these patterns differ from what is observed in *A. thaliana*. In *A. thaliana*, the impact of TEs on adjacent gene expression is extremely consistent across genetically distinct ecotypes originating from the USA (Col-0), Portugal (C24) and Ireland (Bur0; Wang et al. 2013). We thus chose Col-0 as a representative of the species. For *B. distachyon*, less information is available at such a broad geographical scale. We therefore included in our analysis the reference Bd21, which originates from Iraq, as well as ten additional accessions from Spain, France, Iraq and Turkey to further investigate potential within-species variation.

## Results and Discussion

### TEs largely shape genome-wide DNA methylation patterns in Brachypodium

We produced bisulfite- and RNA-sequencing data for eleven *B. distachyon* accessions. SNPs (Gordon et al. 2017; Stritt et al. 2018) and methylation profiling support their split into three distinct genetic clusters (Figure S1): Western (ABR2, Luc1, Bd30-1 and Sig2; Spain and France), Eastern (the reference accession Bd21, Bd21-3, Bd3-1, BdTR12C and Koz3; Iraq and Turkey) and accessions from a third cluster (BdTR7a, Bd1-1; Turkey) showing a delay in flowering time (Gordon et al. 2017; Stritt et al. 2018). In agreement with the genetic distances calculated with SNPs (Gordon et al. 2017; Stritt et al. 2018), the methylation-based PCA confirms that accessions from the Eastern cluster are more closely related to accessions from the Western cluster than to the two other Turkish accessions BdTR7a and Bd1-1. We also used publically available bisulfite- and RNA-sequencing data for the reference accession Col-0 of *A. thaliana* (Table S1).

We first estimated methylation levels (expressed as the percentage of methylated cytosines), gene and TE density in 100 kb windows for the reference genomes of *A. thaliana* (Col-0) and *B. distachyon* (Bd21). The circular visualisation of these three genomic features reveals two distinct architectures: genes and TEs are clearly compartmentalised in *A. thaliana* (Figure 1A) while largely interspersed in *B. distachyon* (Figure 1B). Yet we found that DNA methylation is positively correlated with TE density and negatively correlated with gene density in both species (Figure S2). We then estimated the methylation level of genes and TEs, aggregating the latter at the superfamily level. TE body methylation is significantly higher than gene body methylation in the two species, although variability is observed among TE superfamilies (Figure 1C and D, Table S2-S4 for P-values). Genome-wide DNA methylation patterns are thus largely shaped by TEs in *B. distachyon*, like in other flowering plants species (Zhang et al. 2006; Regulski et al. 2013; Takuno and Gaut 2013; Seymour et al. 2014). We also found that most TE superfamilies and genes show a higher level of body methylation in *B. distachyon* than in *A. thaliana* (Figure 1C and D, Table S5 for P-values), confirming a low genome-wide level of body methylation in *A. thaliana* compared to other species (Seymour et al. 2014, Eichten et al. 2016).

**Figure 1:**
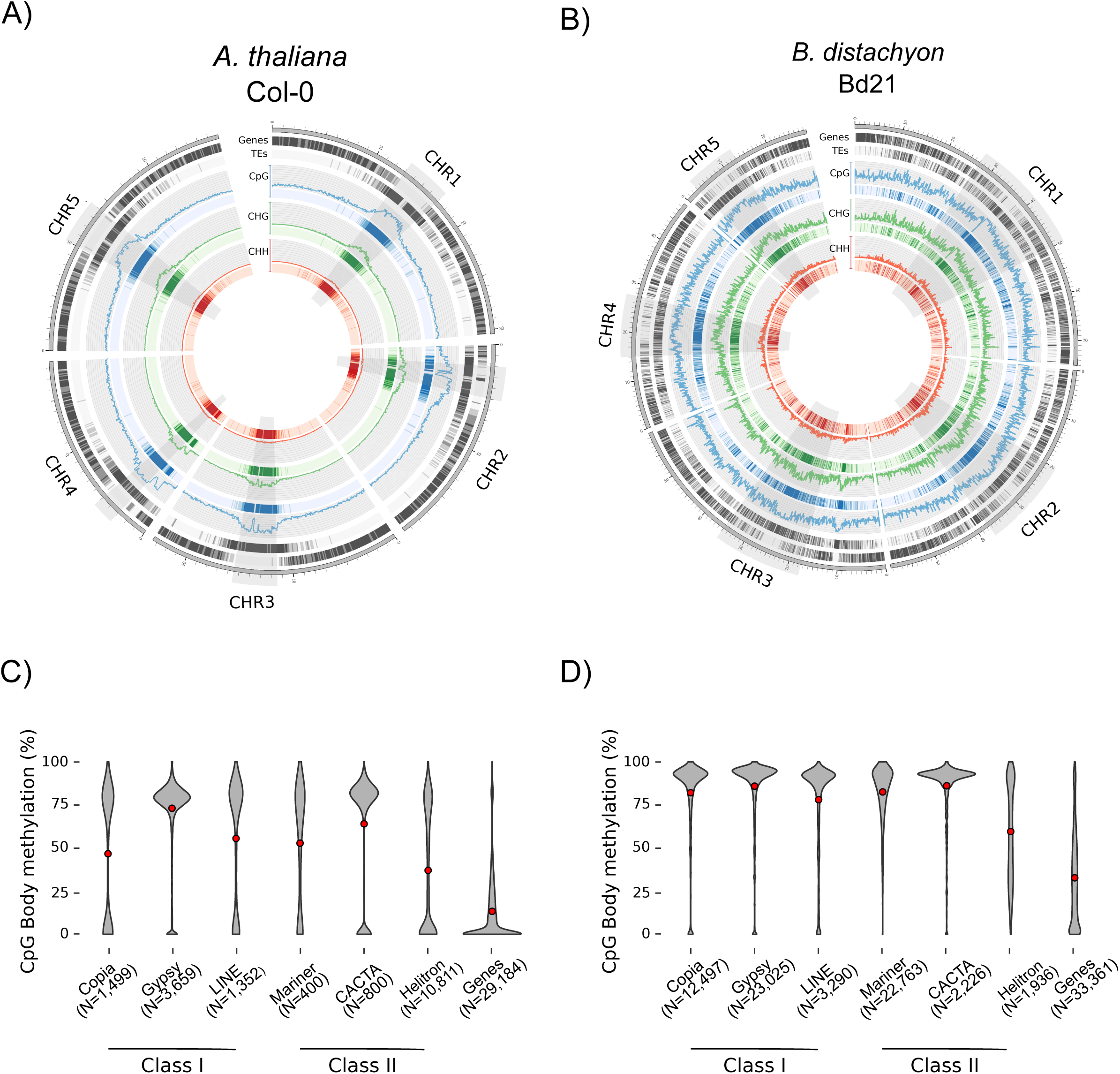
Genome organization and methylation distribution of Col-0 and Bd21. Circos plot displaying methylation levels as well as gene and TE density in 100 kb windows in A) *A. thaliana* and B) *B. distachyon*. Centromeric and pericentromeric regions are highlighted in grey. Panel C and D display CpG methylation levels (expressed as % of methylated cytosines) for the most common TE superfamilies and for genes in *A. thaliana* and *B. distachyon* respectively.

We finally assessed how far methylation spreads around TEs. To do so, we estimated methylation levels in 100 bp windows spanning 2.5 kb around each TE insertion and averaged the values per TE superfamily. We also categorized TE insertions depending on their location on the chromosome (arm vs. centromere) and visualised the results by producing heatmaps. For the three contexts, we observe that DNA methylation around TEs spreads over longer distances in gene-poor centromeric regions than in distal regions of the chromosome arm in *A. thaliana* (Figure 2). In *B. distachyon*, spreading mostly reaches a few 100 bp, except around centromeric *Gypsy* elements where methylation tends to spread beyond 2 kb (Figure 2). Thus, in contrast to our expectations, high levels of TE body methylation in *B. distachyon* are generally not accompanied with a long spreading of methylation, neither in the centromeres nor on the chromosome arms. This is surprising given that extended methylation spreading around TE is associated to gene-poor pericentromeric regions in rice (Choi and Purugganan, 2017), a species closely related to *B. distachyon*. Because patterns are consistent across CpG, CHH and CHG contexts, we concentrated the rest of the analysis on CpG methylation.

**Figure 2:**
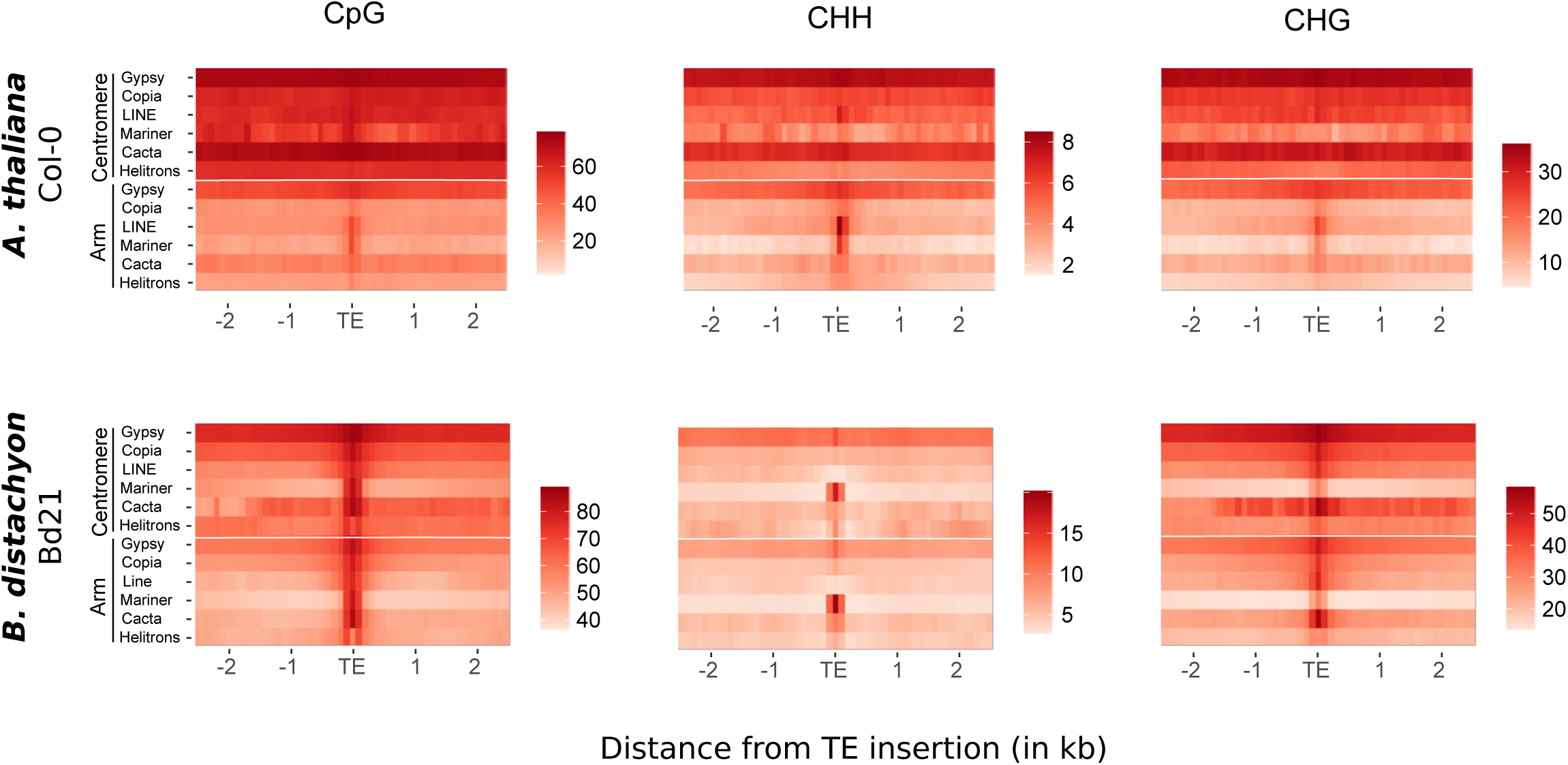
Methylation spreading around transposable elements. Heatmaps representing methylation (CpG) spreading around TEs located in the pericentromere/centromere regions or on the arm of the chromosomes for *A. thaliana* (N=18,521 classified TEs) and *B. distachyon* (N=62,708 classified TEs). Scales were adapted for the six panels to account for the lower genome-wide methylation level in *A. thaliana* and the different methylation levels for the three contexts.

### Impact of TEs on nearby gene expression

In *A. thaliana*, genes flanked by a TE within 2 kb show reduced expression compared to genes devoid of TEs in their vicinity (Wang et al. 2013). It was proposed that this may result from a combined effect of DNA methylation spreading and DNA structure variation (Wang et al. 2013). For both the reference accessions Col-0 and Bd21 we estimated the average level of expression for genes having a TE (regardless of the type of TE) in their sequence or within windows ranging from near vicinity (0-200 bp) to up to 2 kb (Figure S3). While we reproduced the pattern described above with publically available data in *A. thaliana*, our analysis shows that TEs do not have such an effect on nearby gene expression in *B. distachyon*. To rule out potential technical artefacts, we extended the analysis to the ten additional *B. distachyon* accessions we sampled but did not observe any overall effect of TEs on nearby gene expression either (Figure S3). This is also surprising as the distance between a given gene and its closest TE is significantly smaller (Wilcoxon-test, P-value <0.0001; Figure S3) in Bd21 (1,074 bp) than in Col-0 (5,457bp).

The effect of TEs on methylation and gene expression is known to be family-dependent (Eichten et al. 2012; Choi and Purugganan 2017), which might explain why no effect is observed when all TEs are pooled. We therefore repeated this analysis separately for the ten most active TE families (Stritt et al. 2018) in all eleven *B. distachyon* accessions to explore potential genetic background effects. We also produced heatmaps as described above to assess a potential link between methylation spreading around these ten TE families and nearby gene expression. Methylation spreading was in a second step used to statistically cluster the ten families. Eventually, as young TEs might be more efficiently targeted by silencing mechanisms, we extracted the age of each copy from Stritt et al. (2019). For four of the ten families, this analysis allowed to uncover clear effects on gene expression following two distinct patterns (Figure 3A): a short-distance effect (< 400 bp; all P-values <0.001) conveyed by two *Gypsy* families (RLG_Bdis0039 and RLG_Bdis004) and one *Copia* (RLC_Bdis022) family; and a long-distance effect (1 kb; P-values <0.001) for the *Gypsy* element RLG_Bdis180. However, methylation spreading is unlikely to be involved as our hierarchical clustering shows that spreading varies from long (RLG_Bdis004, RLG_Bdis180) to intermediate (RLG_Bdis0039) and short (RLG_Bdis022) around these four families (Figure 3B). In the same manner, there is no clear association neither with the age of the family (Figure 3B) nor with the genetic background as all accessions show comparable gene expression patterns (Figure 3A). It is nonetheless worth noting that RLG_Bdis004 and RLC_Bdis022 are two of the most recently active families in the genome of *B. distachyon* (Stritt et al. 2019), suggesting a possible scenario where *de novo* silencing of these two elements may influence nearby-gene regulation. Altogether, not more than 131 genes are influenced by one of these four families, confirming that the impact of TEs on gene expression is limited in *B. distachyon*.

**Figure 3:**
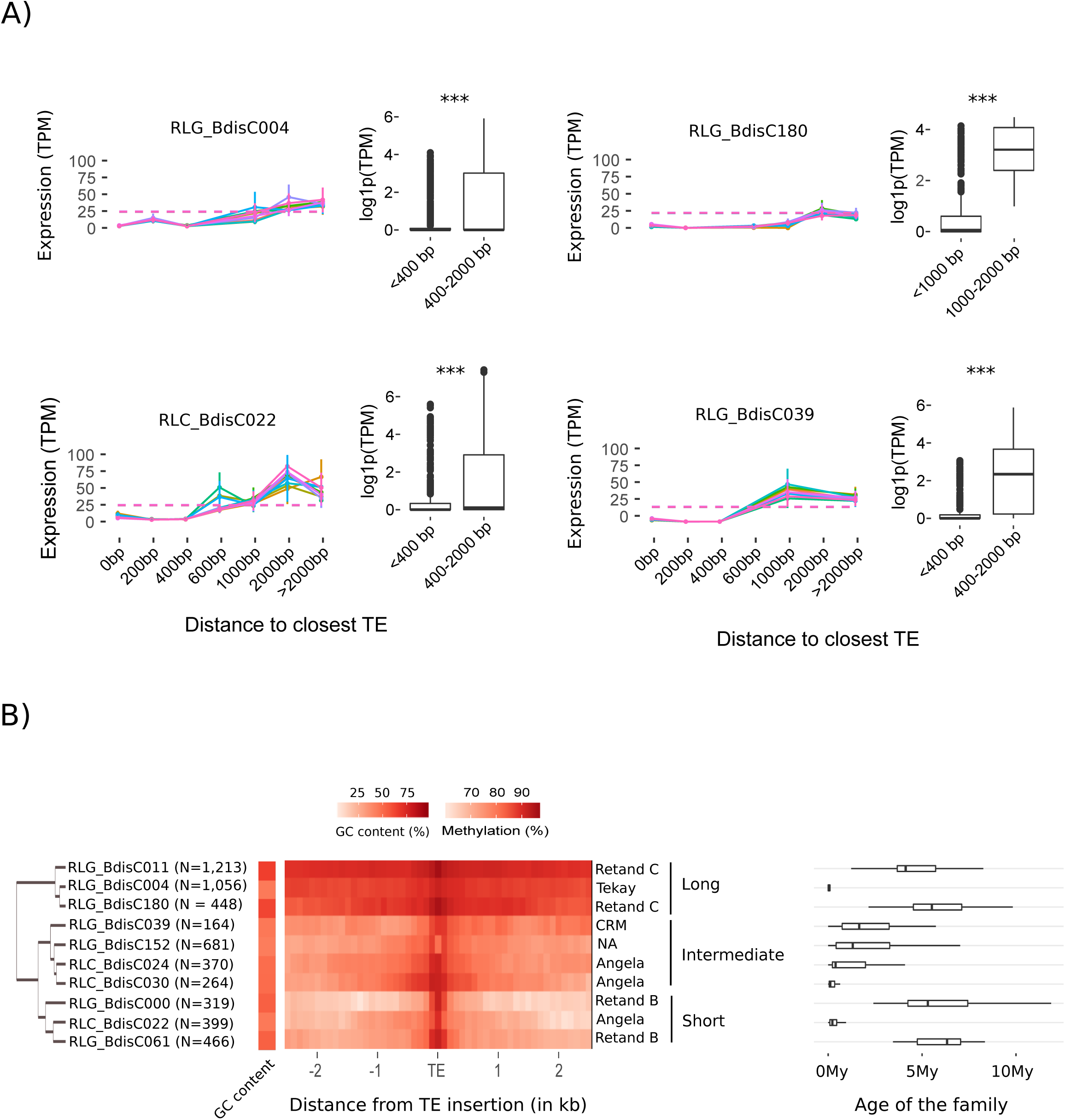
Impact of transposable elements on gene expression. A) Average gene expression as a function of the distance to the nearest TE belonging to the four LTR-retrotransposons RLG_BdisC039, RLG_BdisC004, RLG_BdisC180 and RLC_BdisC022 in the natural accessions of *B. distachyon*. Each coloured line represents a different accession. The dashed line represents the genome-wide level of expression. Boxplots display the expression of genes harbouring a TE in their vicinity for two distance classes only, all accessions pooled together. Stars indicate Wilcoxon test levels of significance: ^***^<0.001 B) Heatmap representing methylation (CpG) spreading around the 10 most active LTR retrotransposon families in Bd21. Families were clustered based on the level of methylation spreading (long, intermediate and short) around each insertion. Names on the right of the heatmap describe the TE lineage to which each family belongs. Boxplots display the age of each families estimated by (Stritt et al. 2019).

We eventually investigated the impact of TE insertion polymorphisms (TIPs) on gene expression. Overall, we identified 1,833 high-confidence TIPs, i.e. insertions present in at least one of the ten natural accessions but absent from the reference genome Bd21 (Table S6). Due to the relative small number of TIPs located nearby genes, we lacked power to investigate their impact on gene expression at the family level. To circumvent this difficulty, we measured the log-fold change in gene expression for loci harbouring a TIP in their vicinity (<1 kb) compared to the same loci without a TIP in a different accession. We found that the majority of the TIPs have no effect on adjacent gene expression (Figure S4), consistent with previous findings that TE polymorphisms have little impact on methylation spreading in *B. distachyon* (Eichten et al. 2016). These results contrast again with what is found in *A. thaliana*, where polymorphic TEs are associated with differential expression of nearby-genes among natural accessions (Stuart et al. 2016). Despite the lack of global effect, a subset of 32 TIPs show a significant effect on nearby gene expression at high confidence (Table S7), providing candidates for further functional validation.

### Genome organization and functional impact of TEs

The differences observed between the two species may result from the nature of their TEs. Notably, the genome of *B. distachyon* display a much higher ratio of solo-LTRs:intact copies for both conserved elements and TIPs (up to 14:1 for some families; Stritt et al. 2019) compared to *A. thaliana*, where the number of intact copies approximate the number of solo-LTRs (Devos et al. 2002). This may reflect a much broader aspect of the TE dynamics in the two species and one could assume that *B. distachyon* harbours overall a much larger proportion of non-functional TEs than *A. thaliana*. As non-functional elements might be less efficiently targeted by silencing, they might have in return a limited influence on their neighbouring genes. This hypothesis, however, does not hold for rice. There, the ratio solo-LTR:intact elements is 1.6:1 (Tian et al. 2009) and therefore does not differ much from the one found in *A. thaliana*. Yet, TEs also have little impact on gene expression in rice (Choi and Purugganan 2017).

As an alternative hypothesis, methylation spreading has been suggested to be under purifying selection (Choi and Purugganan 2017) and such a mechanism may largely constrain the impact of TEs on adjacent genes. Compared to *A. thaliana*, TEs and genes are largely interspersed in both *B. distachyon* and rice (Garg et al. 2015) and selection against the deleterious effect of TEs might be stronger in such systems. We therefore suggest that the differences we observed between *A. thaliana* and *B. distachyon* with regard to the functional impact of TEs are largely inherent to the contrasting architecture of their genomes. In the light of the current results and recent studies (Seymour et al. 2014, Choi and Purugganan 2017, Quesneville 2020), it appears that genome organization varies greatly between *Brassicacea* and grasses. With more than 12,000 species (Christenhusz and Byng 2016), grasses cover about 40% of the planet (Gibson 2009). Generalizing the patterns reported in *A. thaliana* to all plants, as commonly seen in the literature, is therefore arguable. Our study is calling for the analysis of a larger set of plant genomes to reflect the tremendous diversity present in this kingdom.

## Materials and Methods

### Plant material

We used eleven accession of *B. distachyon* originating from Iraq (Bd21, Bd21-3, Bd3-1), Turkey (BdTR12c, BdTR7a, Koz3, Bd1-1) and Spain (ABR2, Sig2, Luc1, Bd30-1). Seeds were sown on soil-sand mixture in 50 ml pots. For bisulfite and for RNA sequencing, three biological replicates per accession were grown. The second and third leaves of each replicate were sampled after 24 days of 16/8 hours of day/night conditions.

### Whole genome bisulfite sequencing (WGBS) and RNA-sequencing analysis

For *B. distachyon*, WGBS libraries were prepared as described in the supplementary material Method S1. For *A. thaliana*, we used publicly available data (Table S1 for references). Raw data were processed as described in Method S1 and mapped onto the reference genomes Bd21 for *B. distachyon* (Phytozome 12, https://phytozome.jgi.doe.gov) and Col-0 for *A. thaliana* (TAIR10, www.arabidopsis.org). Methylation levels were estimated following the method described in (Schultz et al. 2012; Method S1) and by averaging the percentage of methylated cytosines over the three replicates per genotype within each focal unit (e.g. 100 kb/100 bp genomic windows, TE and gene body). We drew Circos plots (Connors et al. 2009) to visualize the architecture of the two reference genomes. The PCA based on methylation data was performed using ggplot2 in R (R development core team 2013). We eventually used GLM models with logit transformation to characterize the relationship between gene or TE density and methylation levels. P-values were obtained with the R package multcomp (Hothorn et al. 2008).

For each of the eleven *B. distachyon* accessions, mRNA was extracted using the RNAeasy Plant Mini Kit from Qiagen. Libraries were prepared by the D-BSSE Basel and paired-end sequencing was performed on a Illumina HiSeq 2500. For *A. thaliana*, we again used publicly available data (Table S1). Reads were mapped to the reference genomes with Salmon quant v.0.12.0 (Patro et al. 2017). The abundance of gene expression was quantified averaging transcripts per million (TPM) per accession for each single genotype over the three replicates.

### Methylation spreading

We estimated methylation spreading by calculating methylation levels in 100 bp windows spanning 2.5 kb upstream and downstream of each TE insertion. TE insertions were categorized with regard to their position in centromeric/pericentromeric regions or on chromosome arms and aggregated at the family-(for the ten most active TE families) or superfamily levels. Heatmaps depicting methylation levels around TEs were drawn in R using ggplot2 (Wickham 2009). We also used the levels of methylation spreading around TEs to calculate Euclidean distances between the ten most active TE families and to perform a hierarchical clustering using hclust in R (R development core team 2013).

### Impact of TEs on gene expression

We assessed the potential link between the presence of a TE in the vicinity of a gene and expression by assigning genes to different categories depending on whether they harbor a TE in their sequence or within windows ranging from near vicinity (0-200 bp) to up to 2 kb. Average gene TPM and standard errors were calculated per distance class with all TEs pooled together as well as for the ten most active families separately and plotted using the R package plotrix (Lemon 2006). For a subset of candidate families, we also compared the expression level (log1p transformed) of genes displaying a TE in their vicinity (0-400 bp for RLG_Bdis0039, RLG_Bdis004 and RLC_Bdis022; 0-1 kb for RLG_Bdis180) to the expression level of genes displaying a TE further away (between 400 bp-2 kb and 1 kb-2 kb respectively) with a Wilcoxon test.

### Transposable element insertion polymorphism analysis

Detection of transposable element insertion polymorphisms (TIP) was performed with detettore (github.com/cstritt/detettore) using sequencing data from (Gordon et al. 2017). Detettore was specifically developed on *B. distachyon*. It allows to output different features, including the presence of target site duplication (TSD), which can be used to filter out false positive. Filtering criteria were adjusted so that no TIP is found when the reads of Bd21 are mapped against the reference genome. The expression of genes harboring a TIP in their vicinity (<1 kb) in at least one of the accessions was compared to the corresponding genes in the other accessions. For each single gene, difference in expression between groups was tested using a t-test with a confidence interval of 0.95. Change in gene expression was considered significant when the P-value was <0.05 and the log2 change >1.

## Acknowledgment

This work was supported by the PSC-syngenta fellowship, the University Research Priority Program Evolution in Action of the University of Zurich and the Swiss National Science Foundation (PZ00P3_154724). The authors would like to thank the D-BSSE for the sequencing effort, the Genetic Diversity Center at ETH Zurich for providing access to their computing resources and Michael Thieme for his comments on the manuscript.

## Data availability

All the data produced for the current study are available under the ENA project number PRJEB39651. For *A. thaliana*, references are provided in the supplementary materials. Scripts are available at https://github.com/mwylerCH.

