## Supplemental Data 1 for "Impact of transposable elements on methylation and gene expression across natural accessions of *Brachypodium distachyon*"

**Supplementary Materials**

**Figure S1: Geographical and genetic origins of the eleven *B. distachyon* accessions.** A) The map displays the country of origin as well the genetic cluster of origin based on SNP data (Gordon et al. 2017, Stritt et al. 2018) B) PCA obtained with the methylation data produced for the eleven accessions, the three replicates pooled together.

**Figure S2: Relationship between gene or transposable element density and methylation levels in Col-0 and Bd21.**

**Figure S3:** **Impact of TEs on gene expression.** Average gene expression as a function of the distance to the nearest TE in *A. thaliana* and in the natural accessions of *B. distachyon,* all TEs pooled together. The table provides the number of TEs for each distance class in the two reference genomes Bd21 and Col-0.

**Figure S4: Transposable element insertion polymorphisms.** Volcano plot displaying expression changes in genes harbouring a TIP in their vicinity. Significant changes are displayed in red.

**Table S1: WGB- and RNA-sequencing output information**

**Table S2: Within-species glm outputs (CpG context)**

**Table S3: Within-species glm outputs (CHG context)**

**Table S4: Within-species glm outputs (CHH context)**

**Table S5: glm outputs - comparison *A. thaliana* vs. *B. distachyon***

**Table S6: Information about the 1,833 TIPs identified across the natural accession of *B. distachyon.***

**Table S7: Information about the 32 high-confidence TIPs**

**Method S1: Extended Materials and Methods**
