## Supplementary material for "Impact of transposable elements on methylation and gene expression across natural accessions of *Brachypodium distachyon*": Method S1

**Methods S1**

*Library preparation for Whole Genome Bisulfite Sequencing*

All *B. distachyon* samples were processed as described in (Stritt et al. 2019). Shortly, leaf samples where disrupted with Bead Ruptor 24 (OMNI). DNA was extracted with the DNeasy Plant Mini Kit (Quiagen) and shared with a Q800R2 Sonicator (Qsonica). Libraries were prepared with the Kapa Hyper Prep Kit (Kapa Biosystems) using methylated NimbleGen SeqCap adapters (Roche) and bisulfite converted with EZ DNA Methylation-Gold (ZYMO Research). Bisulfite converted libraries were PCR amplified with five PCR cycles and the subsequent products were size selected using AMPure XP Beads (Agencourt). Quality and amount of all prepared libraries were assessed using a TapeStation 2200 (Agilent) and a Real Time qPCR, respectively. Sequencing was performed on an Illumina HiSeq 2500 (Illumina) with 151 paired-end reads.

*Whole Genome Bisulfite Sequencing processing*

Processing, alignment and extraction of cytosine methylation levels was performed as described previously in (Stritt et al. 2019). Adaptor sequence were trimmed using trim_galore (0.4.5, bioinformatics.babraham.ac.uk/projects/trim_galore). The remaining reads were mapped with Bismark (0.19.0, Krueger & Andrews, 2011) using Bowtie2 (2.3.2, Langmead & Salzberg, 2012) to the reference genomes Bd21 (v.3.0, Phytozome 12, https://phytozome.jgi.doe.gov/pz/portal.html) and Col-0 (TAIR10, www.arabidopsis.org) for *B. distachyon* and *A. thaliana*, respectively. Deduplication was performed using the deduplicate_bismark script from Bismark. Subsequently, the methylation level for each single cytosine was obtained for the three CpG, CHG and CHH context with the Bismark script bismark_methylation_extractor with --comprehensive --bedGraph --CX --ignore 2 --ignore_r2 1 settings. Single conversion efficiencies were calculated using the unmethylated chloroplast sequence.

*RNA-sequencing analysis*

For each of the eleven *B. distachyon* accessions, mRNA was extracted using the RNeasy Plant Mini Kit from Qiagen. Library preparation and paired-end sequencing (2x100 bp) were performed by the Genomics Facility Basel (ETH Zurich) on a Illumina HiSeq 2500. For *B. distachyon* and for *A. thaliana*, the CDS of Bd21 (v.3.1, Phytozome 12, https://phytozome.jgi.doe.gov/pz/portal.html) and *Col-0* (TAIR 10, www.arabidopsis.org) were used as references. Reads were mapped with Salmon quant (version 0.12.0, (Patro et al. 2017) with automatic library type detection. The abundance of gene expression was quantified averaging the Transcripts Per Million per accession for each single genotype over the three replicates.
