## Supplementary figures and images for "Impact of transposable elements on methylation and gene expression across natural accessions of *Brachypodium distachyon*"

### Supplementary Figures

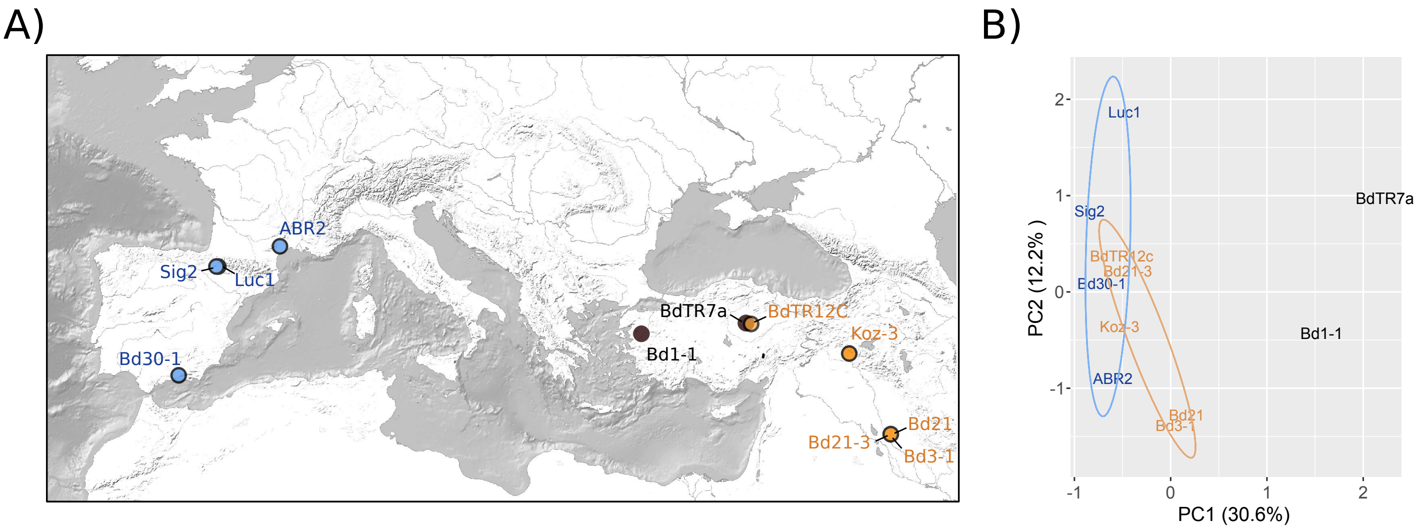

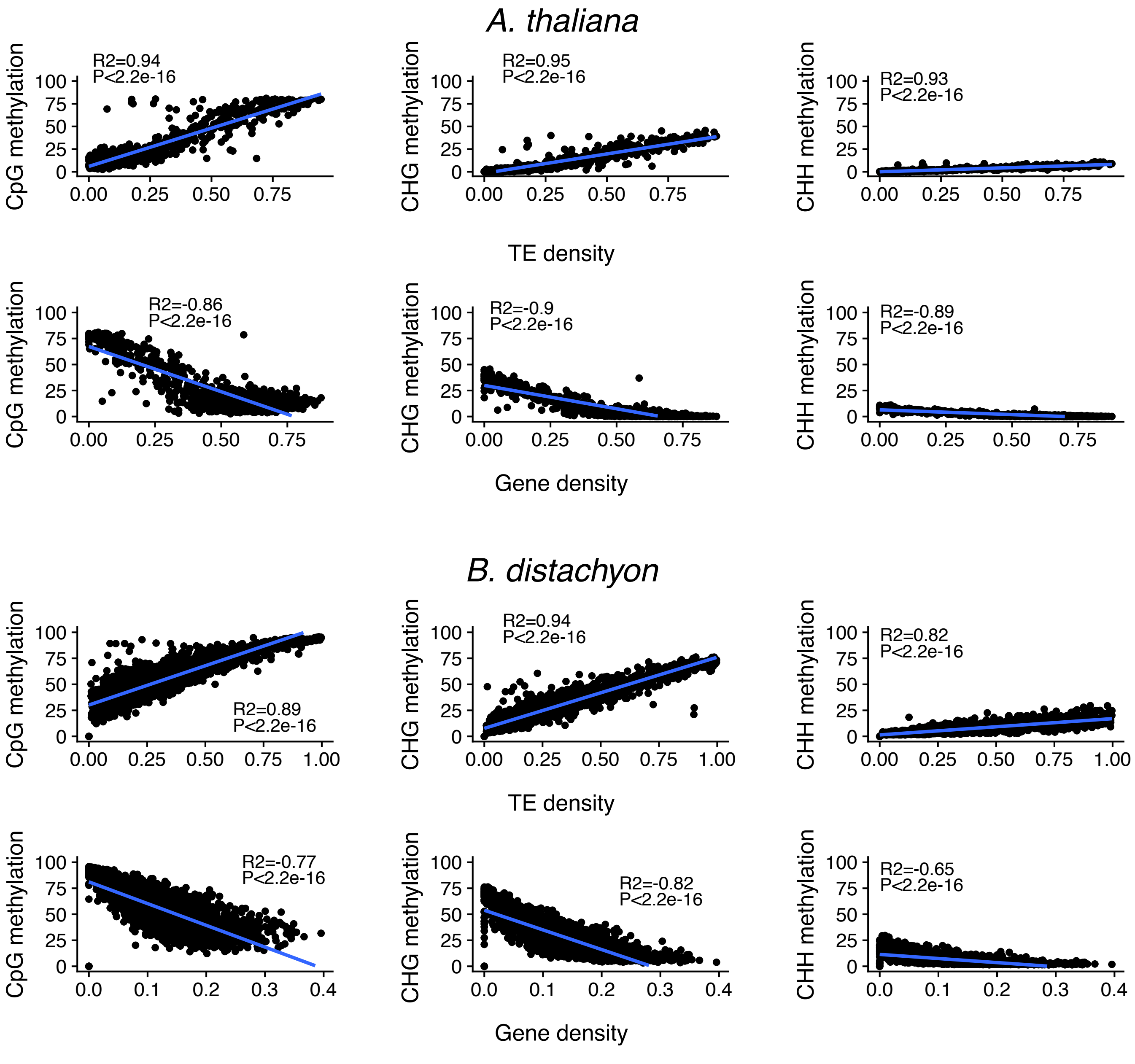

**Figure S3**

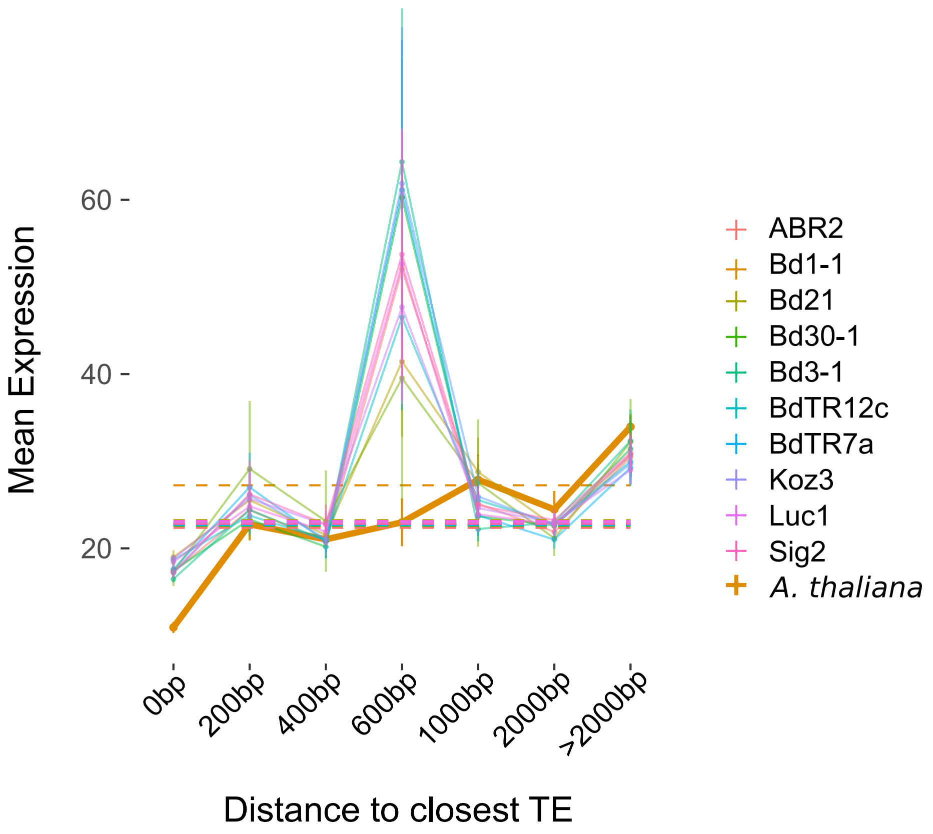


| | Bd21 | Col-0 |
|---|---|---|
| within gene | 21877 | 5285 |
| 0-200bp | 2273 | 3862 |
| 200-400bp | 2197 | 2127 |
| 400-600bp | 1757 | 1362 |
| 600-1000bp | 2591 | 2008 |
| 1000-2000bp | 3651 | 3368 |
| >2000bp | 6408 | 19547 |

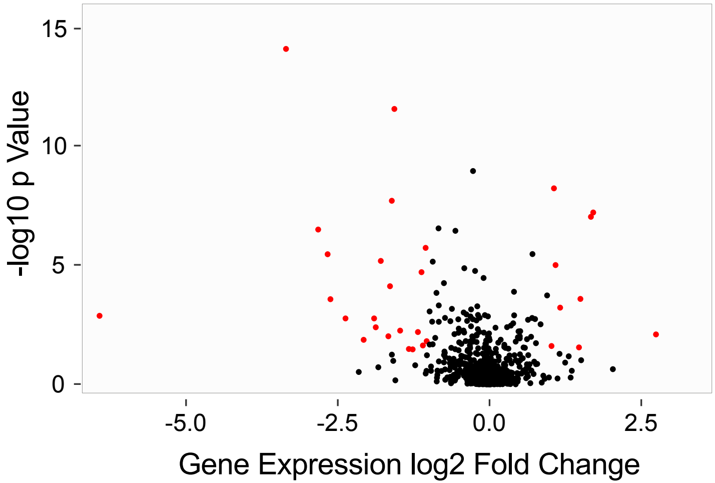
